# Macrophage-mediated immunoediting drives ductal carcinoma evolution: Space is the game changer

**DOI:** 10.1101/594598

**Authors:** Chandler Gatenbee, Jeffrey West, Annie M. Baker, Nafia Guljar, Louise Jones, Trevor A. Graham, Mark Robertson-Tessi, Alexander R. A. Anderson

## Abstract

Under normal conditions, the immune system is capable of rapidly detecting and eliminating potentially dangerous entities, including tumor cells. Due to intense selection pressure imposed by the immune response, tumor cells often evolve strategies to avoid elimination in a process known as immunoediting. It is less known how the evolutionary response to immune predation is altered by context. We explore the evolution of immune escape strategies in ductal cancers, a natural case in which to study evolution in different contexts: inside and outside of ducts. We highlight the role of macrophages as a source of “public goods,” releasing diffusible factors (reactive oxygen species and growth factors). Immunohistochemistry reveals differences between macrophage densities of invasive ductal carcinomas and non-invasive ductal carcinomas *in situ*. For the first time, immunohistochemistry (IHC) imaging data comparing DCIS to IDC were used to initialize mechanistic agent-based models of evolutionary dynamics. By using IHC to map the initial conditions of a growing tumor, we show that spatial competition and structure influence transient dynamics during invasion. These dynamics are context-dependent, a conclusion that may be missed from interpreting imaging or non-spatial modeling alone. Before invasion, the presence of macrophages correlate with shorter ductal breach times. After invasion, tumors may employ a “pioneer-engineer” strategy where pioneering immunoresistant cells on the tumor’s edge stimulate the release of M1-macrophage-derived reactive oxygen species, degrading surrounding stroma. Behind the invasive edge, the engineering immunosuppressive cells promote the release of M2-macrophage-derived growth factors, providing a long-term immune escape strategy. Together, mathematical modeling and image analysis highlight the crucial role tumor-associated macrophages play in immune escape and invasion, both inside and outside of ducts.

## 1 Introduction

Tumor cells stimulate immune attack by the expression of tumor specific neoantigens, danger associated molecular patterns (DAMPs), stress ligands, and lack of major histocompatibility complex (MHC) class I molecules, among other mechanisms^1, 2^. The immune response is complex; here we provide a brief overview of tumor-immune interactions (see Figure 1). The immune response is initiated after antigen presenting cells, such as dendritic cells (DCs), encounter the tumor and then migrate to secondary lymphoid organs. There they present antigens to T-cells (both CD4^+^ and CD8^+^) and plasma cells^3–7^. Antigen-specific immune cells will be activated, go through rapid expansion, and produce inflammatory cytokines. This is followed by an influx of cytotoxic CD8^+^ T-cells, Type 1 and Type 2 helper T-cells (Th1 and Th2, respectively), regulatory T-cells (Tregs), macrophages, and B-cells to the site of the tumor. Cytotoxic T-cells, Th1 cells, and classically activated macrophages (M1) attack the tumor cells carrying the cognate antigen^3, 8, 9^. Additionally, M1 macrophages release reactive oxygen species (ROS), potentially inducing apoptosis in surrounding cells, while natural killer (NK) cells may destroy cells that do not express MHC class I proteins^3, 9–12^. Once the threat clears, the immunosuppressive cytokine interleukin 10 (IL-10) is released by cytotoxic T and Th1 cells, beginning the process of down-regulating inflammation^9, 13^. In addition to IL-10, Tregs release TGF-*β*, a powerful anti-inflammatory cytokine that inhibits antigen presentation by DCs, the activity of IFN-*γ*, the proliferation of T-cells, and the cytotoxic activity of NK and T cells^14–16^. IL-10 and IL-4 produced by Th2 and Tregs also push the macrophage phenotype from an inflammatory M1 state to the pro-growth and wound-repair M2 phenotype that releases growth factors, matrix metalloproteinases (MMPs), and vascular endothelial growth factor (VEGF), all of which aid in tissue repair and restoration of homeostasis^13, 17^.

**Figure 1.**
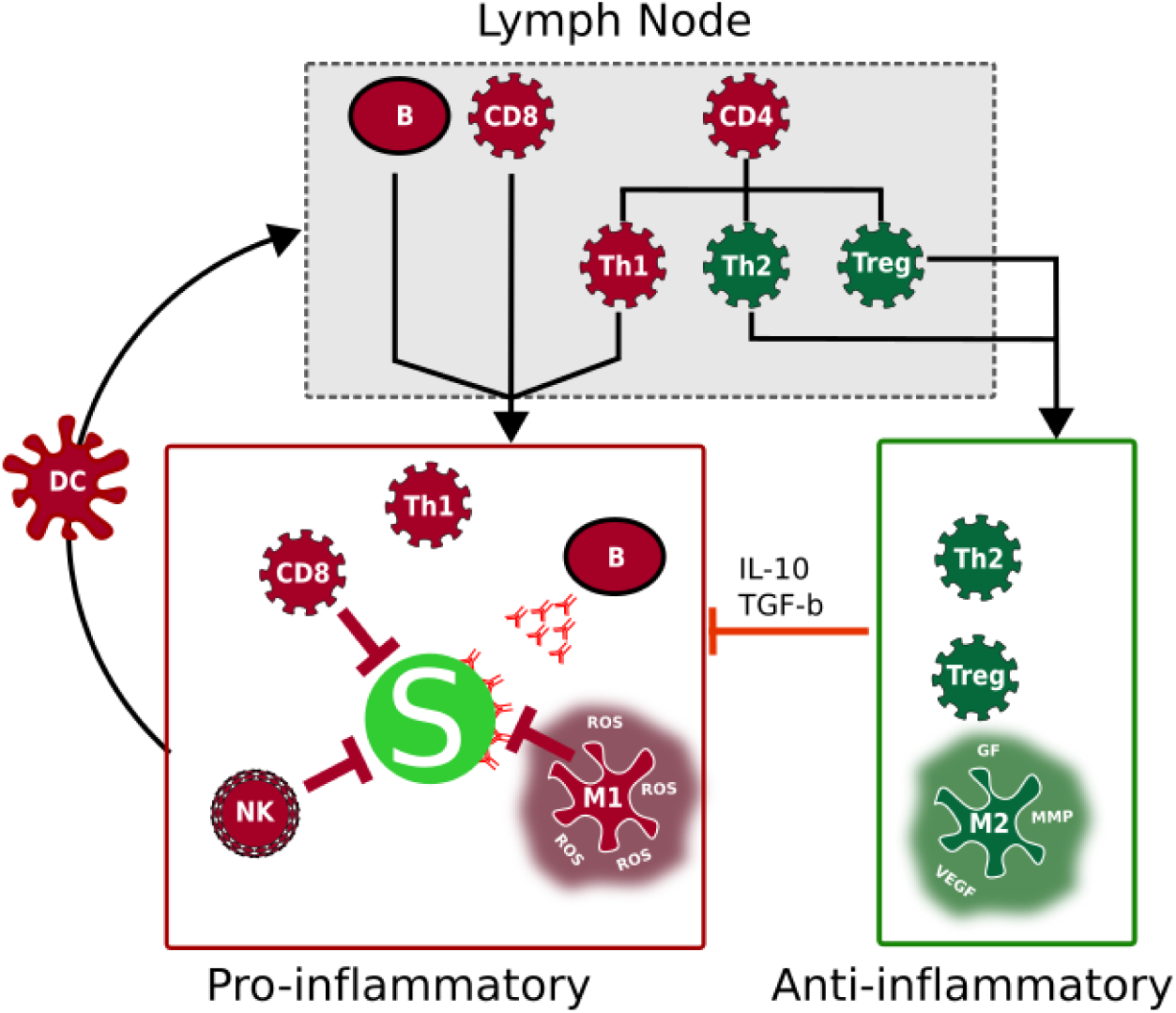
Normal immune response to cancer. Tumor cells (*S*) elicit immune attack due to the expression of antigens and production of a variety of signals. The immune response is initiated when DC present antigens to immune cells in lymph nodes. Pro-inflammatory immune cells migrate to the site of the tumor and attempt to remove it. Eventually, anti-inflammatory cells reduce inflammation and release signals to repair damage caused during tumor attack.

Immune response against tumors imposes strong selection pressures that favor cancer cells that have evolved strategies to escape elimination^5, 15, 24^. This immunoediting process occurs in three phases: elimination, equilibrium, and escape. Here, we focus on immune escape, in which tumor cell variants that have evolved the capacity to avoid elimination now have the opportunity to grow unchecked by the immune response^5, 15^. Tumors employ various immune escape strategies which we classify into three broad categories: immunoevasion, immunoresistance, and immunosuppression (Figure 2). Immunoevasive strategies (E) allow tumor cells to avoid detection by the immune system, often due to defective antigen presentation pathways or even a lack of any detectable antigens. Immunoresistant strategies (R) encompass mechanisms that protect the cell from elimination, despite having elicited an inflammatory immune response. An example of this strategy is the expression of checkpoint inhibitors, such as PD-L1, which prevent cells from being eliminated by cytotoxic T-cells^20^. Immunosuppressive strategies (I) give cells the ability to down-regulate the anti-tumor immune response, often by recruiting other immune cells (e.g. M2 macrophages, Tregs, and neutrophils) that release anti-inflammatory cytokines. A more comprehensive list mapping mechanisms to immune escape strategies can found in Table 1, and see^5, 11, 18, 25–27^ for in depth reviews.

**Table 1.** Defining Immune Escape Strategies

| Strategy | Mechanism | Example | Ref. |
| --- | --- | --- | --- |
| Resistant | Prevent apoptosis at level of receptors | Loss of TRAIL | 18 |
|  |  | Loss of Fas (CD95) | 11,18 |
|  |  | Loss of MICA or MICB | 11,15 |
|  |  | Increased expression of CD47 | 19 |
|  |  | PD-L1 | 20 |
|  | Internal inhibition of apoptosis | Overexpression FLIP | 14 |
|  |  | Overexpression of Bcl-2 family of proteins | 5,18 |
|  |  | Activation of of Survivin | 14,18 |
|  |  | Activation of cIAP2 | 11,14,18 |
|  |  | Inactivation of FADD | 11 |
|  |  | Inactivation of caspase-10 | 11 |
|  |  | Expression of MUC1 | 14 |
|  |  | Loss of Apaf1 | 14,18 |
|  | Blocking of cytotoxic signals | SPI-6 | 11,18 |
|  |  | PI-9 | 14 |
| Evasive | Loss/decreased expression of MHC-I | Loss of HLA locus on chromosome 6 | 11 |
|  |  | Over-expression of c-Myc | 11 |
|  | Impaired antigen presentation | Inactivation of LMP-2 and LMP-7 | 18 |
|  |  | Inactivation TAP-1 and TAP-2 | 11,14 |
|  |  | Down-regulation of HLA-1 and HLA-2 | 11 |
|  | Evade NK | Inactivation of Necl-2 | 14 |
|  |  | Loss of B7-1 or B7-2 expression | 11 |
|  |  | Loss of CD40 | 11 |
| Suppressive | Diffusible factors | VEGF | 14,15 |
|  |  | IL-10 | 11,14 |
|  |  | PGE2 | 11,14 |
| | | TGF-β | 15 |
|  |  | IDO | 5,14,21 |
|  |  | Phosphatidylserine | 15 |
|  |  | Galectin | 5 |
|  |  | MUC1 | 14 |
|  |  | COX-2 | 14 |
|  | Immunosuppressive immune cell recruitment | Tregs | 5 |
|  |  | MDSCs | 5 |
|  |  | M2 macrophages | 17,22 |
|  |  | Neutrophils | 23 |

**Figure 2.**
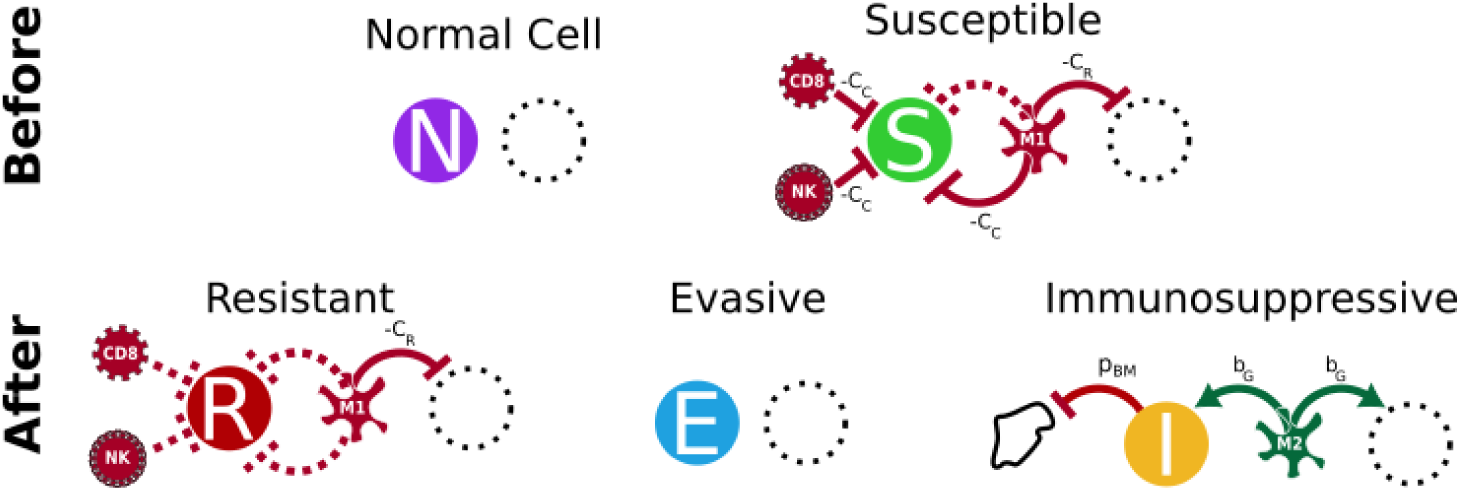
Designing an evolutionary game of immune escape. Top row: before immune detection. Bottom row: immune escape strategies, after immune detection. Each strategy is assumed to attract a complement of immune cells: Susceptible cells (*S*) and immunoresistant (*R*) cells stimulate inflammatory immune cells (M1 macrophages, CD8+ T cells, and NK cells); immunoevasive (*E*) and normal (*N*) cells do not attract immune cells; and immunosuppressive cells (*I*) promote M2 macrophages. Red lines indicate the presence of a cost, while green lines represent benefits. Dotted lines indicate costs and benefits that have been avoided by the strategy. Dotted circles represent neighboring cells that receive bystander costs and benefits of interacting with cells of each strategy. The parameters are defined as: *b*_*G*_ = benefit of binding diffusible growth factors; *b*_*N*_ = increased fitness of tumor relative to normal; *c*_*C*_ = cost of contact with anti-tumor immune cells; *c*_*R*_ = cost of absorbing ROS; *c*_*I*_ = inherent cost of immune escape strategy.

### 1.1 Ductal carcinomas as a case of changing context

Cells using adaptive Darwinian strategies compete within a “context” – the heterogeneous local microenvironment of beneficial growth factors, harmful ROS, stromal cells, immune cells, and other tumor cells. A change in this context (e.g. ductal carcinomas becoming invasive) alters the fundamental nature of competition within the tumor. For example, niche construction (whereby a tumor cell modifies the local microenvironment to boost its fitness relative to neighboring tissue) is important in ductal invasion^28^ and metastasis^29^. Tumor subpopulations can be defined by adaptive “strategies”, which are context-dependent: influenced by neighbor interactions and tumor microenvironment factors rather than the intrinsic molecular properties of the cell alone^28^.

Here we examine ductal cancers in general (breast cancer in particular) as they provide a natural case in which context may influence tumor evolutionary dynamics. Inside the duct, competition is primarily between tumor cells alone, while post-invasion competition transitions to one in which tumor cells also compete with normal tissue. We use evolutionary game theory (EGT) to model immune escape and examine how the selection pressures imposed by the different intra-and extra-ductal microenvironments shape invasion dynamics and tumor heterogeneity. We then use immunohistochemistry (IHC) to quantify the density of intraductal macrophages, the source of public goods, in ductal carcinoma in situ (DCIS) and invasive ductal carcinoma (IDC) and determine if, as predicted by the model, the density of macrophages (and thus the density of public ‘goods’ and ‘bads’) is associated with invasion.

Previous mathematical models designed to study cooperative or competitive interactions between tumor and immune cells have quantified the proliferation response of the immune system to changing tumor burden ^30–32^ and response to therapy ^33^. Others have used agent-based models ^34^ or hematoxylin and eosin–stained images ^35^ to show the importance of spatial organization of heterogeneous immune-evasive “hotspots” within a tumor, especially in the context of competition for shared resources (e.g. during glycolysis). We employ a comprehensive modeling and analysis strategy to predict the evolutionary response of immune escape by combining non-spatial game theory, agent-based models, and image analysis. This provides a complete picture of the role of macrophages as a source of public good in pre-invasive ductal carcinomas.

### 1.2 Evolution dynamics of immune response

Evolutionary game theory is a modeling technique that is ideal to study complex competitive cellular interactions. Many such games have been used to model the interactions between tumor cells and the cells in their microenvironment^36–41^, evolving tumor cell strategies during tumor progression^42, 43^ and treatment dynamics^44–46^. We now introduce and analyze an EGT model based on the replicator system^47^ and extend the same payoff table to a spatial model to show the emergence of spatially-dependent mechanisms of immune escape and invasion.

In this evolutionary game, the “players” are tumor and normal stromal cells. A fitness landscape defined by the payoff table and the replicator equation (see Methods) controls the competition and evolution of these five cell types: normal healthy tissue (*N*); tumor cells that are susceptible (*S*) to detection and elimination by the immune response; tumor cells with the evasive strategy (*E*) that are not detected by the immune system and thus do not elicit a response; tumor cells with the resistant strategy (*R*) which still express antigens and stimulate an immune response but are not eliminated during attack; and tumor cells with the immunosuppression strategy (*I*), which shift the anti-tumor inflammatory response to one that promotes cellular growth and tissue repair.

While cytotoxic T-cells and NK cells usually kill tumor cells through targeted cell-cell interactions, it is noteworthy that macrophages produce diffusible factors that have off-target effects on neighboring cells. This feature of macrophages makes them a source of public “goods” or public “bads” in the game (see^48^ for a public goods game), producing growth factors and DNA damage-inducing ROS, respectively. Thus, each immune escape strategy (evasion, resistance, and suppression) utilizes different mechanisms to gain advantage on a fitness landscape mediated by the public resources produced by macrophages (Figure 2). The model does not explicitly consider immune cells as agents (e.g. M1/M2 macrophages, CD8^+^ T cells, NK cells) but rather models them as downstream costs and benefits arising from immune interactions as detailed in figure 2. Inside the payoff table, *A*, there are three cost parameters (*c*_*I*_, *c*_*C*_, *c*_*R*_ *>* 0) and two benefit parameters (*b*_*G*_, *b*_*N*_ *>* 0). Each row corresponds to the payoff that a cell type receives over each competing cell type (column).

### 1.3 Immune escape as an evolutionary game

Conceptually, the payoff table *A* (see Methods) is best understood by first considering cancer initiation before immune attack (Figure 2, top row). Immune-susceptible tumor cells (S) have a fitness advantage over normal tissue (N), indicated by *b*_*N*_ in the S row, N column (i.e. a ‘driver’ mutation). In the absence of immune response, susceptible cells will grow at the expense of surrounding normal tissue. After immune response is initiated, susceptible cells, which have no immune-escape mechanism, are targeted by T-cells, NK cells, and M1 macrophages, reflected in the parameter *c*_*C*_ in row S.

The three immune escape strategies (*E, I, R*, Figure 2, bottom row) develop mechanisms to evade this immune killing, but this induces an associated immune escape cost (*c*_*I*_) for maintaining this mechanism (added to rows *E, I*, and *R*). We assume that the cost of susceptibility to immune attack, *c*_*C*_, must be greater than the cost of immune strategy, *c*_*I*_, otherwise no immune escape strategies would evolve. Immunosuppressive cells attract M2 macrophages that release public goods in the form of growth factors into the microenvironment. This confers a benefit *b*_*G*_ to cells that interact with immunosuppressive cells, which is added to the *I* row and *I* column of.

Healthy tissue does not stimulate an immune response, so neither cost (*c*_*C*_, *c*_*I*_) is included in row *N*. Resistant cells do stimulate an inflammatory response. Since M1 macrophages release ROS into the microenvironment, cells that cannot process ROS incur a cost, *c*_*R*_, when interacting with either susceptible or resistant tumor cells (added in row *N*, columns *R* and *S*). In this sense, M1 macrophages are distributing the public bad of ROS in the game. We assume that the cost of absorbing ROS is not conferred to tumor cells because tumor cells often express antioxidant proteins to detoxify from ROS^49^.

### 1.4 Initialization of spatial evolutionary game from multi-color immunohistochemistry

This non-spatial model of immune response (replicator equation, see Methods) is extended to include space and structure by implementing an agent based model (ABM) wherein each agent’s fitness is determined by the cost and benefits of interacting with cells in its local Moore neighborhood. The game is played out on a regular lattice where at each time step cells calculate their fitness relative to the surrounding neighborhood of cells (see Methods for more details).

The spatial model simulations are initialized with a patient-specific duct “mask,” *D*, derived from image analysis (see Section 4.3) to define the initial position of tumor cells in the ABM (see Figure 4A). A random mix of tumor strategies (i.e. all strategies except *N*) are placed within the ducts, while only normal cells exist outside of the duct. The border of the duct is composed of a basement membrane (BM) that keeps tumor cells contained within the duct. Since M2 macrophages are capable of producing MMPs that can breakdown the BM of ducts^17, 50, 51^, only cells with strategy *I* are capable of breaking out of the duct during model simulations. A cell with strategy *I* can work to break down neighboring BM (modeled as a small probability of BM destruction, *p*_*BM*_ = 0.01, see Figure 2). Following breakdown of the BM, any cell can divide into that now empty space, simulating invasion from the duct into normal tissue.

## 2 Results

We present results from the non-spatial model (Figure 3), the spatially-explicit agent-based model (Figure 4), and analysis from IHC imaging data (Figure 6), examining the evolutionary response to immune attack in both pre-and post-breach ductal carcinomas. Importantly, the combined analysis of all three approaches reveals the role of macrophages as the source of a public good in a two-phase, context-dependent evolutionary adaptation to immune predation.

**Figure 3.**
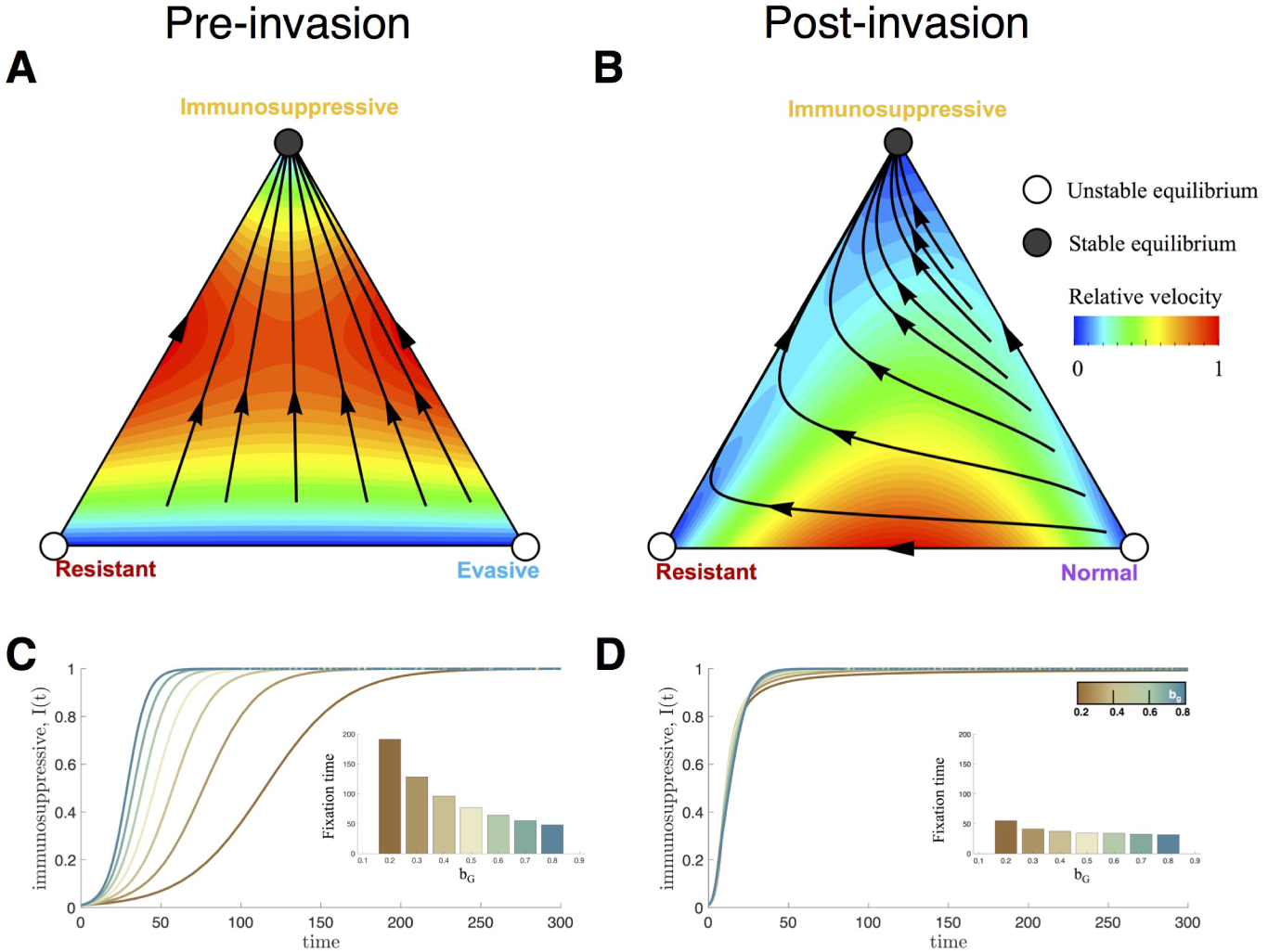
Immunosuppressive strategies dominate before and after ductal invasion. Trilinear coordinate phase portraits are shown with sample trajectories (black lines, see Methods, Non-spatial replicator dynamics; Equations (1) and (2)), color-coded by relative velocity. Circles indicate unstable (open circle) and stable (closed) equilibria. (a) Phase portrait of the three highest fitness players for pre-invasion dynamics before the duct is breached (*b*_*G*_ = 0.5, *b*_*N*_ = 0.5, *c*_*C*_ = 1, *c*_*I*_ = 0.2, *c*_*R*_ = 0.5). All internal trajectories lead to domination of the immunosuppressive strategy. (b) Phase portrait of the three highest fitness players in post-invasion dynamics. With the addition of normal cells, the resistant strategy has a temporary advantage over normal cells (right-to-left dynamics), even though *I* remains the global stable equilibrium. (c,d) The presence of macrophages (i.e. increased *b*_*g*_) pre-(c) and post-invasion (d) leads to a global steady state with complete fixation of strategy *I*. Insets show that the time to fixation is exponentially longer with a linear decrease in *b*_*g*_ inside the duct, yet relatively unchanged outside the duct.

**Figure 4.**
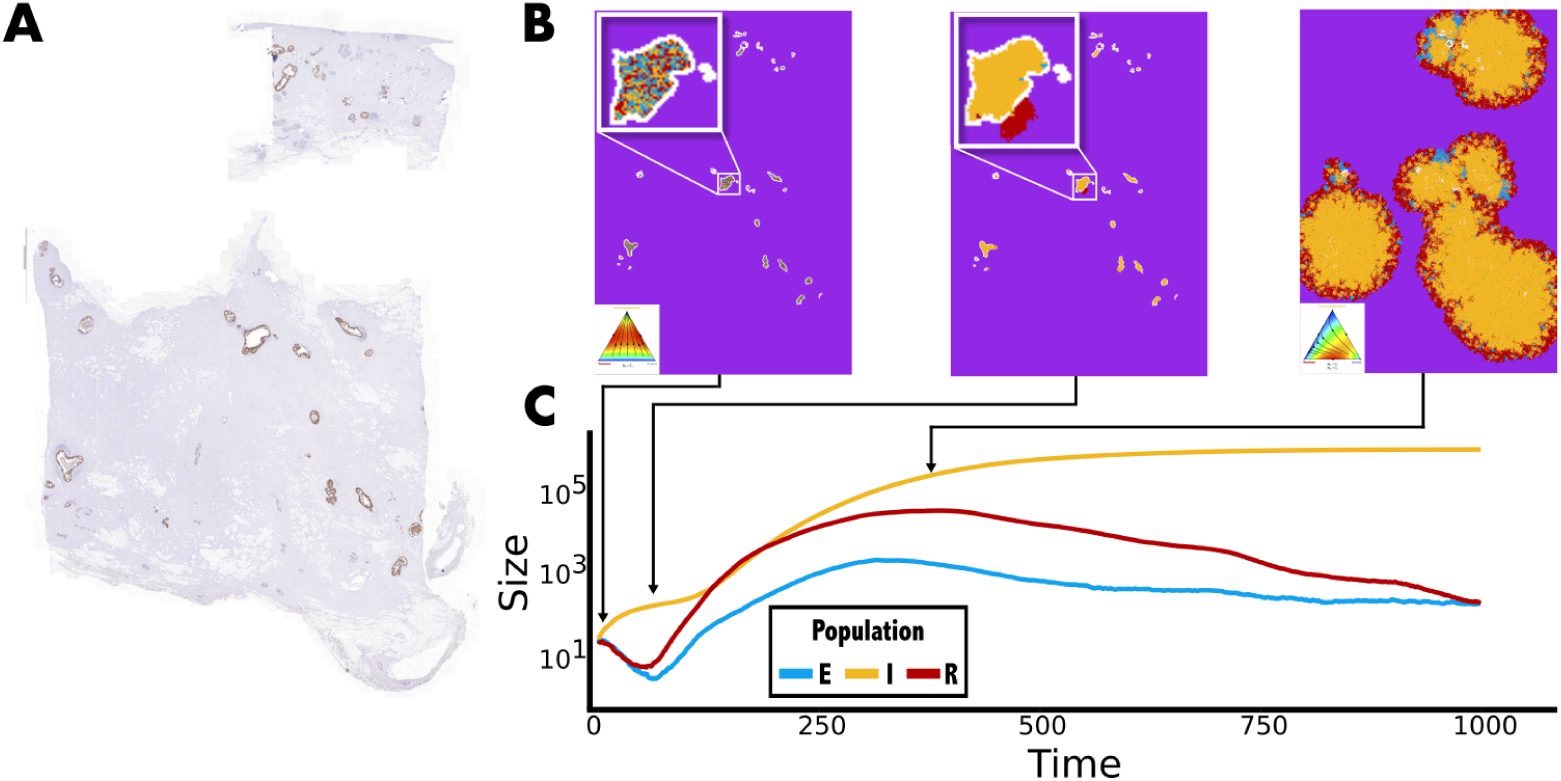
Spatial game of immune escape in ductal cancers seeded by immunohistochemistry. A) The spatial model was initialized using a DCIS sample stained for HER2 (brown). Each pixel positive for HER2 was considered to be a tumor cell, and was assigned a random strategy. B) Snapshots from example simulation. Top left: each duct, outlined in white, is initialized with a random mix of escape strategies. Top middle: The immunosuppressive strategy *I* wins inside the duct and eventually breaches the BM. This is followed by invasion of the immunoresistant strategy *R* into normal tissue (purple). Top right: *R* drives invasion on the tumor edge, while *I* comprises the bulk of the tumor core. C) temporal dynamics of the spatial game, beginning with the intra-ductal dynamics, followed by invasion dynamics. The arrows correspond to the snapshot of the simulations in panel B. Purple=normal cells, blue=evasive strategy, yellow=immunosuppressive strategy, and red=resistant strategy. (*b*_*G*_ = 0.5, *b*_*N*_ = 0.5, *c*_*C*_ = 1, *c*_*I*_ = 0.2, *c*_*R*_ = 0.5)

### 2.1 Analysis of costs and benefits: public goods and public bads

We begin by analyzing competition between strategies inside the duct. Since normal cells are not present there, we ignore competition with strategy *N* for the moment. In the pairwise interactions between *S* and each immune-escape strategy (*E, I*, and *R*), it is clear that the fitness *f*_*i*_ of strategy *S* is always less than the fitness of each immune-escape strategy for all non-zero values of the susceptible population, *x*_4_. The three immune-escape strategies are therefore the dominant players within the duct, and we visualize these strategies on a simplex (Figure 3A), directly from Eq. (1). The corners of this trilinear coordinate frame represent the saturation of a single cell type (e.g. the top corner represents 100% immunosuppressive cells in the population), and the edges indicate pairwise competition.

For the case where cells are constrained within the duct (pre-invasion, Figure 3, left column), the pairwise interactions are shown on the edges: 1) *I* wins over *E*, 2) *I* wins over *R*, and 3) *E* and *R* are neutral. The interior of the triangles contain sample dynamic trajectories (black lines) derived using replicator dynamics. The background color within the triangle indicates the instantaneous relative velocity along the trajectories^52^. All internal initial conditions eventually lead to a stable equilibrium consisting of all *I*.

We now explore how the dynamics change *after* invasion into the normal tissue (*N*). From the payoff matrix, we can see that rows *E* and *R* are identical, except for their interactions with normal tissue through the parameter *c*_*R*_ (see the *N* row, where the resistant strategy *R* has the advantage of releasing ROS and thereby decreasing the fitness of *N*). Thus the three dominant players outside the duct are *N, R*, and *I*, shown in Figure 3, right column. Note that for invasion of *I* and *R* to be possible, we additionally make the respective assumptions that *b*_*G*_ > *c*_*I*_, *b*_*N*_ > *c*_*I*_, and *c*_*R*_ > *c*_*I*_. Again, the pairwise interactions are shown on the edges: 1) *I* wins over *R*, 2) *I* wins over *N*, and 3) *R* wins over *N*. Interestingly, the resistant cell type shows a *transient* competitive edge over the normal population, before eventual saturation by the immunosuppressive population. Generally, evolutionary games are analyzed for long-term evolutionarily stable states (ESS), while transients are sometimes ignored. Here we have an important transient dynamic, in that the strategy *R* appears to invade into normal tissue faster than *I*, even as *I* eventually dominates over *R*.

It is reasonable to assume there will be variation in the densities of macrophages that are recruited by *I*. We model this variation by changing *b*_*G*_, the parameter that represents the benefit macrophages provide to the immunosuppressive strategy. An increased benefit (due to increased ductal density of macrophages) leads to a decrease in fixation time of *I* within the duct (Figure 3C), but a negligible difference in post-invasion fixation (Figure 3D). This suggests that the increased presence of macrophages in the duct may be correlated with a shorter time to a membrane breach, while having little impact on post-breach dynamics.

### 2.2 Emergence of spatially self-assorting pioneer and engineer phenotypes

In order to explore the importance of ductal structure and thus intra-and extra-ductal contexts, we now introduce a spatial version of the game. The simulation is initialized using a slice of tissue taken from a DCIS biopsy that had been stained for HER2 (see Section 4.3 for a description of the method to segment HER2). Normally, cells are not located within ducts, and so each pixel within the duct that is positive for HER2 is assumed to be associated with a tumor cell. By using immunohistochemistry to map the initial conditions of a growing tumor, we can get a more realistic idea of how space and structure influence invasion dynamics, both inside and outside of ducts.

The general dynamics of the non-spatial model are recapitulated in the spatial model (see Figure 4). When the players are all tumor cells (e.g. *E, I, R, S*) without population *N*, strategy *I* quickly sweeps to fixation. Outside the duct, where *N* is now in the game, *R* has a transient advantage, while *I* gradually takes over.

However, while the overall dynamics display similar trends, the spatial model highlights the important role *R* plays after invasion. As seen in Figure 4B, *R* drives invasion through the normal tissue as an edge layer in the tumor mass, while *I* takes over from the core, replacing the leading *R* cells as they expand. This pattern of “pioneer” phenotypes on the edge of the tumor which invest in invasive strategies that permit them to acquire new resources and “engineer” phenotypes in the tumor core that aggressively compete for limited resources has been previously noted in breast cancer, but from a metabolic perspective^53^. Here, this similar pattern emerges because *R* stimulates the release of M1-macrophage-derived ROS, lowering the fitness of *N* on the tumor edge. Conversely, *I* releases M2-derived growth factors, which increases the fitness of *N*; for this reason, *I* is not as invasive as *R*, while still dominating the tumor in the long term. It is also worth noting that the spatial organization allows the evasive strategy *E* to compete in small pockets (see Figure 4, blue). This is a localized effect, occurring only when *E* and *R* are competing with *I* in the absence of adjacent normal tissue. The simulation depicted in Figure 4 is also shown in supplemental video 1.

### 2.3 A context-dependent two-phase public goods game

The role of context-dependent interactions is more clear: although the immunosuppressive strategy is able to facilitate invasion by breaking down ductal walls, the rate of tumor growth is highly coupled with the presence of normal and resistant cells. Figure 5 compares the non-spatial model with the spatial model, with and without the presence of the immunoresistant cell population. After invasion, the tumor’s strategy of coupling a pioneer (*R*) with an engineer (*I*) facilitates rapid tumor growth compared with a singular immune escape strategy; *I* alone grows more than twice as slowly as the combination.

**Figure 5.**
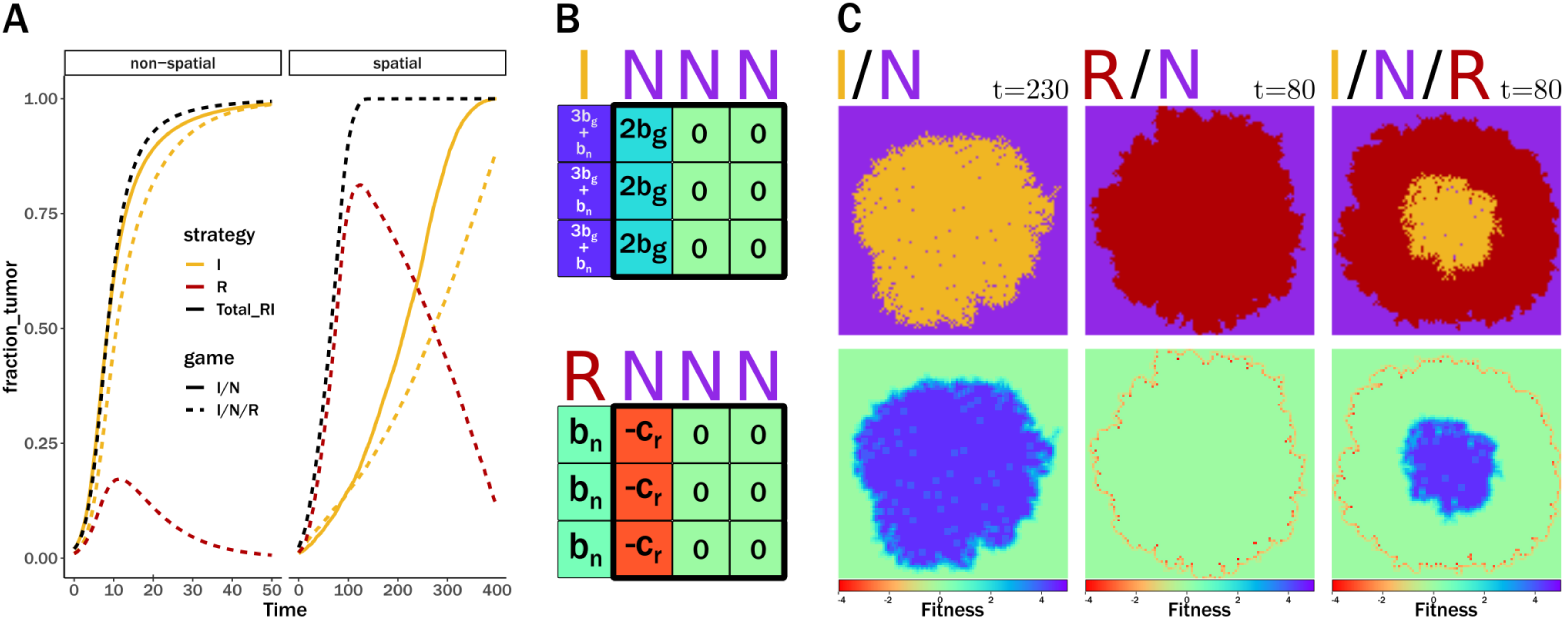
Emergence of an engineer-pioneer effect in the spatial game. A) Population dynamics in various games played out in the normal tissue, following ductal breach. *I*/*N* is the game in which the *I* strategy competes only with *N* (solid lines), normal tissue, and *I*/*N*/*R* is the game where the *I, N*, and *R* strategies are all present (dashed lines). In the spatial game, the importance of the transient increase in *R* is clear. *R* plays the role of pioneer, dramatically increasing the rate of tumor growth (black lines) compared to when only *I* is present (yellow lines). In contrast, the impact of this transient response is minimal in the non-spatial game, as all populations interact with each other and there is no opportunity to be a pioneer. B) In the non-spatial game, all cells interact with one another, and thus there is no invasive front. However, in the spatial game, the death/birth process that occurs at the invasive edge dictates the rate of tumor growth. In the model, a “death box” (bold black line) is placed around a random cell, and the cell with the lowest fitness is removed. If this box is at the *I*/*N* interface, then the least fit cells (*f* = 0) will never be adjacent to the *I* population, and therefore it will be replaced by a normal cell. However, if the box lands at the *R*/*N* interface, the least fit cells (*f* =*-c_R_*) will always be adjacent to *R*, which will then proliferate into the available space. This causes rapid invasion by *R* into normal tissue, increasing the rate of tumor growth. Shifting the death box one position to the left has the same result. C) Visualizing fitness in the spatial game. Due to the death/birth process, normal cells at the invasive front of *I* have a higher fitness then typical *N* cells in the population, slowing tumor growth. Conversely, normal cells at the invasive front of *R* have the lowest fitness (red ring), allowing *R* to rapidly invade.

This pioneer and engineer relationship is only resolved in the spatially explicitly model: the pioneer resistant cells self-organize on the tumor’s invasive outer edge. Without considering both context (invasion into normal tissue) and spatial dependence (self-organization of tumor strategies) of tumor immunoevasion, immune-escape strategies may be falsely concluded to be interchangeable (see Figure 5, left).

### 2.4 Imaging analysis supports link between macrophage density and invasiveness

Figure 6 compares the density of macrophages observed in histological samples of DCIS and IDC, using sections from formalin-fixed paraffin-embedded blocks (5 cases of pure ductal carcinoma in situ and 5 cases of invasive ductal carcinoma). Comparisons of DAB stains for CD68 positivity between DCIS and IDC ducts give a measure of the density of macrophages.

**Figure 6.**
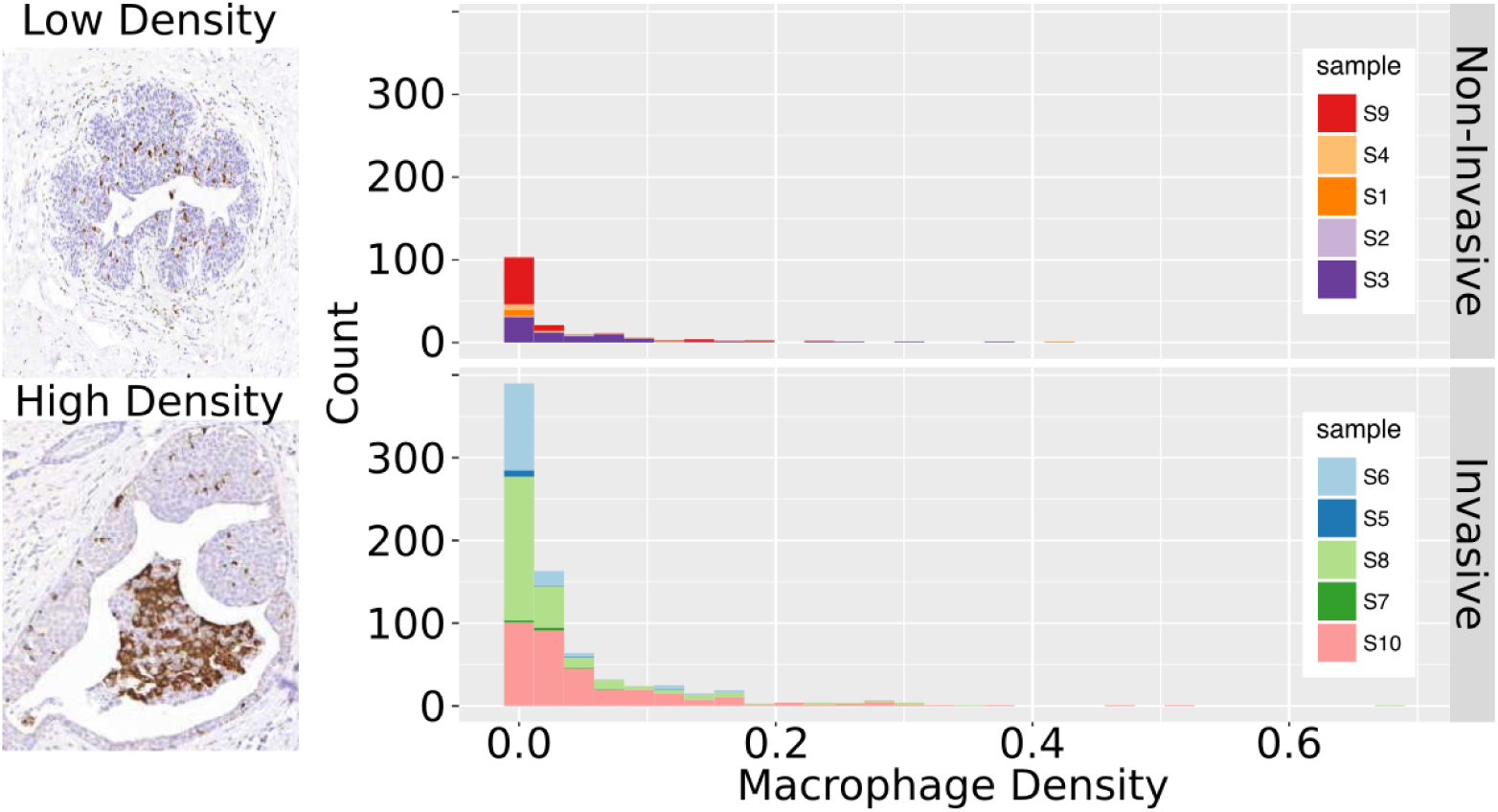
invasive ductal carcinomas have increased intra-ductal macrophage density. The intra-ductal density of macrophages was determined for samples with pure DCIS and those with IDC using immunohistochemistry. Examples of intra-ductal densities can be seen on the left. Samples with IDC had significantly higher densities of macrophages than those with pure DCIS (Wilcox Rank Sum test (*p <* 0.001), Kolmogorov–Smirnov (*p <* 0.001)).

Two representative ducts are shown, one of a DCIS duct (top left) with low macrophage density (brown) and an IDC duct (bottom left) with a high macrophage density. For each duct, the macrophage density is calculated by the ratio of DAB-positive pixels to non-transparent pixels within the duct, shown as a histogram in figure 6 (see Methods for more details). These data indicate that IDC samples have higher intra-ductal densities of macrophages than DCIS samples. This was confirmed with significance via two statistical tests: the Wilcox Rank Sum test, a non-parametric test that compares two sample distributions; and the Kolmogorov–Smirnov (K-S) test, a non-parametric test that compares empirical cumulative distribution functions (ECDF). The Wilcox Rank Sum test value was *W* = 75293 (*p <* 0.001). The K-S test had a value of *D* = 0.22463 (*p <* 0.001). A lower ECDF indicates there are fewer ducts with low macrophage densities, and thus that more ducts in IDC have higher macrophage densities than in IDC. While there have been studies of describing the connection between poor prognosis and macrophage density in the stroma and surrounding ducts^54, 55^, we are unaware of other studies that have quantified the intra-ductal macrophage density. Post-invasion, there is some evidence of ROS-stress in tumor adjacent stroma, which damages surrounding healthy tissue in prostate cancer^56^.

## 3 Discussion

Results from both the spatial and non-spatial model emphasize the importance of context in shaping evolutionary dynamics (see figure 7). In the first phase (pre-invasion), macrophages are the source of the “public good” that drives immune escape dynamics. The immunosuppressive population growth dominates but is highly sensitive to macrophage density, and begins to break down ductal walls (figure 7, top middle). In the second phase, the introduction of interactions with normal tissue following duct breach facilitates the transient survival of the less fit immunoresistant (*R*) strategy on the tumor edges (figure 7, top right). The pioneering immunoresistant population facilitates faster invasion of the engineer, the immunosuppressive population. The ROS produced by the associated M1 macrophages accelerates invasion, regardless of M2 macrophage density. This important dynamic emerges only when the environmental differences inside and outside of the duct are considered, namely the competition with normal cells in the stroma (figure 7, bottom, dashed lines). This result also emphasizes that cell fate is not entirely determined by intrinsic factors, but that spatially-determined context plays a critical role in shaping evolutionary dynamics.

**Figure 7.**
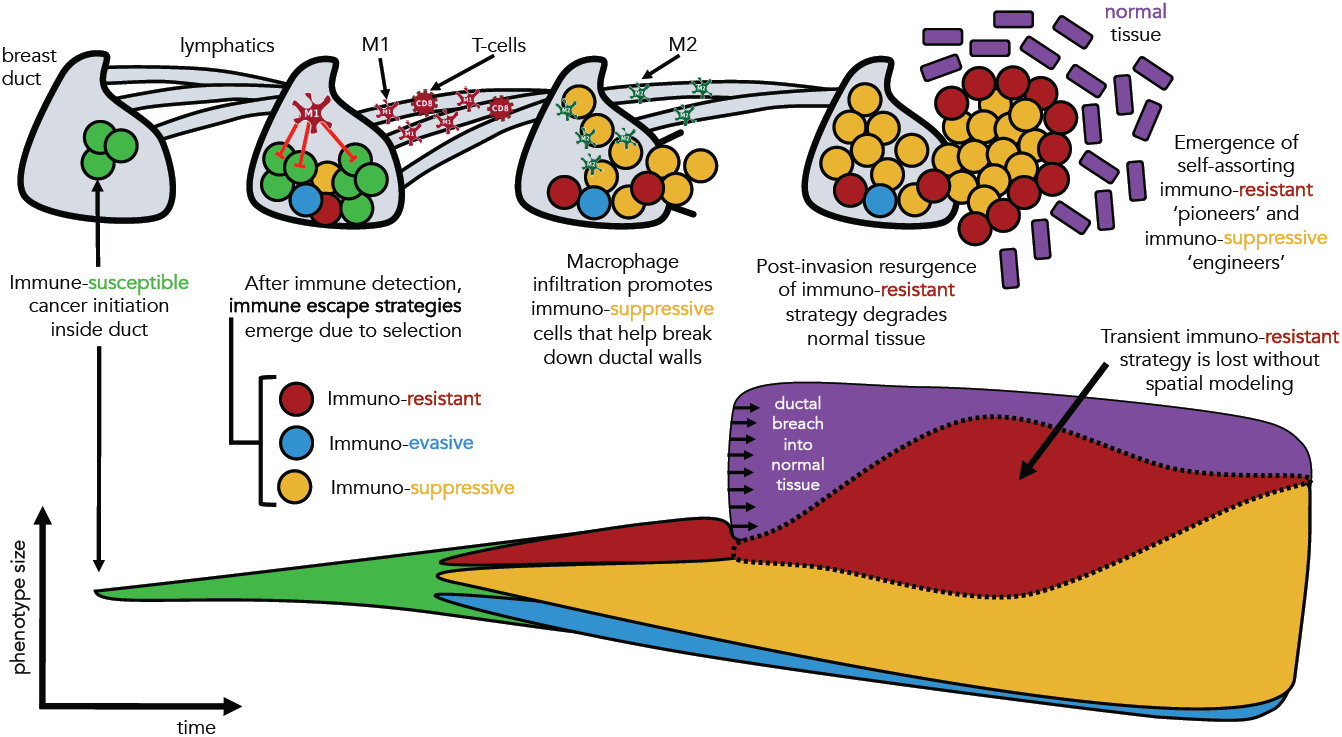
Summary schematic. Cancer initiation in the breast ducts (green cells) may be susceptible to immune attack. Upon immune detection, several immune escape strategies emerge due to selection. Immuno-suppressive strategies emerge as the dominant strategy, facilitating ductal breach. After breaching the duct, immuno-resistant strategies form a self-assorting ‘pioneering’ edge, degrading normal tissue. This effect would be lost without spatial modeling, indicated by the dashed line in the phenotype abundance schematic (bottom).

In the model, cells with an immunosuppressive strategy sweep to fixation within the duct, which in turn increases the density of the recruited M2 macrophages. Since M2 macrophages release MMPs that facilitate breaching of the duct wall, a prediction of the model is that macrophage density should be associated with an increased risk of invasion. Quantification of intra-ductal macrophage density using IHC with CD68 as a macrophage marker does suggest that tissues with invasive tumors, such as IDC, do indeed have higher densities of intra-ductal macrophages than do tissues without invasion, like DCIS.

Comparison of the results from the spatial and non-spatial models reveals how the explicit modeling of space and structure alters the evolutionary dynamics within a tumor. In this case, the significance of the transient increase in frequency of the resistant strategy *R* is only apparent in the spatial model due to the self-assortment of the “pioneering” resistant cell type. In the non-spatial model, the importance of the transiently-increased *R* frequency is less clear, while in the spatial model it is readily apparent that the increase in *R* is due to its ability to invade the stroma more rapidly than any other strategy. Modeling space also emphasizes the importance of population structures that may emerge. Here, the co-existence of *R* with *I*, which is only possible with spatial segregation of *I* in the tumor core and *R* at the tumor’s invasive edge, results in a more aggressive tumor. By including space, the model leads to important insights that were obfuscated in the non-spatial model.

Results from this study provide mechanistic insight into the evolutionary dynamics of immune-evasive strategies at play during the transition from DCIS to IDC. The potential role of macrophages in mediating this transition, and the correlation of their ductal densities and invasiveness emerges both from image analysis and the evolutionary competition mechanisms posited in the mathematical models. Our results suggest that patient-specific intra-ductal macrophage density may be a prognostic marker of invasion as well as a mechanism for increased tumor heterogeneity, and that these mechanisms must be analyzed within a spatial context.

## 4 Methods

In this evolutionary game theory model the “players” are tumor and normal healthy tissue. A fitness landscape defined by the payoff table and the replicator equation (see equation 3) controls competition between these five cell types: immunosuppressive tumor cells (*I*), immunoevasive tumor cells (*E*), immunoresistant tumor cells (*R*), susceptible tumor cells (*S*), and normal healthy tissue (*N*). Macrophages produce diffusing factors modelled as public “goods” (growth factors) or public “bads” (reactive oxygen species). Interactions between cells employing various immune escape strategies and their neighbors are mediated by these public goods and bads.

### 4.1 Non-spatial replicator dynamics

The model is based on a five compartment replicator system of equations, which is a deterministic birth-death process in which birth and death rates are functions of cell fitness, and cell fitness is a function of prevalence in the population. The vector of cell population fractions 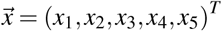 is modeled using Eq. (1), where *i* = 1, 2, 3, 4, 5 corresponds to *E, I, R, S*, and *N* cells, respectively. The individual population fitness, *f*_*i*_, is given in Eq. (2), and is compared to the average fitness of all five populations, given by (*f*) = ∑_*i*_ *f*_*i*_*x*_*i*_.

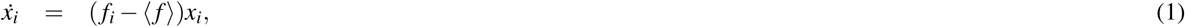

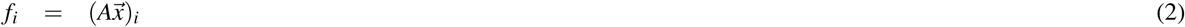

If the fitness of sub-population *i* is greater than the average fitness (i.e. *f*_*i*_*–⟨f⟩ >* 0), that sub-population grows exponentially, whereas if it is less (*f*_*i*_ –*⟨f⟩<* 0), it decays. The fitness is a function of the payoff matrix *A* that includes the effect of public goods and bads, shown in Eq. (3).

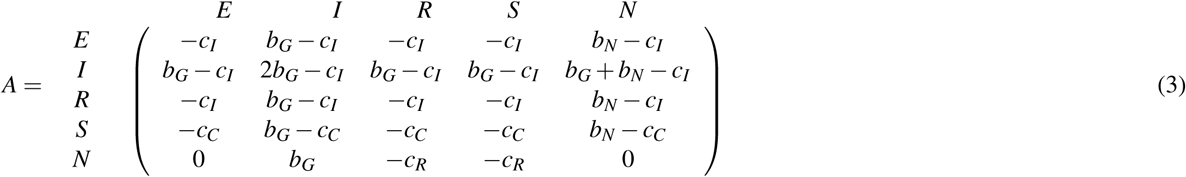

### 4.2 Spatial dynamics

A spatially-explicit extension of the evolutionary game is played out on a regular square lattice where at each time step cells calculate their fitness relative to the surrounding neighborhood of cells (e.g. *E, I, R, S*, or *N*), and then using the payoff matrix *A* to sum the total costs and benefits of interacting with those neighbors. That is, the fitness of a given cell with strategy *i* is *f*_*i*_ = ∑_*k*_ *A*_*ik*_, *k ∈ M*, where *M* is the set of the cell’s neighbors, each with strategy *k*. The ABM algorithm randomly selects *n* focal cells during each time step. For each focal cell, the least fit cell in the cell’s Moore neighborhood is removed (chosen stochastically if multiple cells share the lowest fitness value), and the fittest cell from the removed cell’s neighborhood produces a daughter in the empty position. Finally, dividing tumor cells may undergo mutation with probability *µ* = 0.01, producing a cell with a different immune escape strategy chosen at random from the four tumor cell types.

### 4.3 Quantifying ductal macrophage density with immunohistochemistry

Multi-color immunohistochemistry images were analyzed to estimate macrophage density and create ductal masks for ten cases. Sections of formalin-fixed paraffin-embedded blocks (5 cases of pure ductal carcinoma in situ (DCIS); 5 cases of invasive ductal carcinoma (IDC)) were deparaffinised and rehydrated, before blocking endogenous peroxidase activity with 3% H_2_O_2_ for 20 minutes at room temperature (RT). Antigen retrieval was performed at 95°C, pH 6.0 for 20 minutes before cooling to RT. Sections were blocked in 2% goat serum and 1% bovine serum albumin for 1 hour at RT, before incubation with mouse anti-human CD68 antibody at 1:100 dilution (M0876, Dako UK Ltd, Ely) for 1 hour at RT. Signal amplification was performed by incubating with biotinylated anti-mouse secondary antibody (E0354, Dako) at 1:400 dilution for 30 minutes at RT, and Streptavidin-HRP (P0397, Dako) at 1:500 dilution for 30 minutes at RT. Detection was performed by incubating with 3,3’-diaminobenzidine (DAB) for 90 seconds, and sections were lightly counterstained with Gill’s haematoxylin before dehydration and mounting.

Comparisons of DAB stains for CD68 positivity between DCIS and IDC ducts give a measure of the density of macrophages. A mask, *D*, covering only unbreached ducts was created (using Python OpenCV^57^) and DAB was segmented from the image by converting the RGB image to the LUV colorspace and selecting only those pixels with 0 <*L* <200 and 145 <*V* <255. The number of transparent pixels in each duct was determined by converting the RGB image to optical density space, and then counting the number of pixels that have a value less than 0.15 for all three channels, as described in^58^.

For each duct *i*, the macrophage density is calculated by 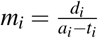, where *d*_*i*_ is the number of pixels positive for DAB, *a*_*i*_ is the total number of pixels within duct *i*, and *ti* is the total number of transparent pixels within duct *i*.This ductal mask is an *i*.initial condition for the ABM simulation to show the transient dynamics during invasion.

